# Genetic ancestry-specific meQTLs control immune function regulation in a breast cancer cohort of African and European patients

**DOI:** 10.1101/2024.08.29.610316

**Authors:** Kyriaki Founta, Nyasha Chambwe

## Abstract

We explored the impact of genetic ancestry on DNA methylation (DNAm) in African and European ancestry breast cancer patients. Analyzing data from 578 subjects in The Cancer Genome Atlas (TCGA) cohort, we identified 757 differentially methylated sites associated with genetic ancestry (aDMS). Methylation quantitative trait loci (meQTL) mapping showed that the majority of aDMS are regulated by multiple SNPs differentially prevalent by ancestry. Notably, expression quantitative methylation (eQTM) mapping linked 95% of aDMS to local gene expression, indicating potential impact on gene regulation.

Furthermore, 72% of ancestry eQTMs were regulated by meQTLs, underscoring the genetic regulation of DNAm variation affecting the transcriptome. Immune response regulation, particularly involving HLA genes, was the most enriched process among ancestry eQTMs. Additionally, 31% of differentially expressed genes between groups defined by genetic ancestry were under ancestry eQTM control. These findings highlight the gene regulatory potential of ancestry-associated DNAm and its importance in population-specific analyses.

## Introduction

Black women of African descent face an excessive burden of breast cancer both globally and in the United States1. They have a higher incidence of the most aggressive and invasive breast cancer subtype, triple-negative breast cancer (TNBC), and worse TNBC-associated clinical outcomes compared to other population groups^2–4^. These racial and ethnic cancer health disparities (CHDs), resulting in unequal cancer burden and poorer outcomes across different populations, remain an under-addressed issue and a persistent clinical challenge^5,6^.

Although previous research has explored the impact of genetic factors associated with African ancestry^7,8^ and the implications of socioeconomic factors such as healthcare and treatment access on breast cancer disparities^6,9^, the molecular basis of the more aggressive biological and clinical behavior of breast cancer in Black women of African ancestry is not yet fully understood. Elevated signals of homologous recombination deficiency in breast cancer^7,8^ and alterations in key cancer genes across various cancer types^10^ have been linked to African ancestry. Additionally, specific gene expression markers have been observed in breast tumors of African ancestry, with some of these markers also detected in normal African breast tissues^4^. Social Determinants of Health (SDoH) have been also associated with breast cancer gene expression alterations^4^, and interactions between environmental factors and ancestry-specific mutations have been reported for other cancer types^10^. However, the exact mechanisms through which these complex and inter-related factors interact remain unclear.

DNA methylation (DNAm), a crucial epigenetic modification in development, helps establish and maintain cellular identity in mammals^11^ while serving as a link between genetics and environmental exposures^12^. DNAm levels are regulated by individual genotypes at specific loci, known as methylation quantitative trait loci (meQTLs)^13^, but they can also change in response to developmental or environmental cues^11^. Several studies have investigated population-specific DNAm, suggesting these differences may act as evolutionary mediators between genetic and phenotypic variation^14,15^. The impact of genetic variation and population-specific alleles on DNAm specificity has been previously reported; multiple studies identifying ancestry-related meQTLs have highlighted the involvement of the genetic architecture of DNAm in nuclear regulatory pathways^16^ and their potential implications in numerous phenotypic traits^17^.

Dysregulated DNAm has been associated with cancer, with global hypomethylation observed in cancer tissues and site-specific hypermethylation leading to the epigenetic silencing of tumor suppressor genes^18^. Aberrant DNAm is widely observed in breast cancer^19,20^. Although genetic ancestry-specific DNAm patterns have been identified among breast cancer patients^21^, most studies examining DNAm in the context of CHDs consider DNAm differences exclusively as markers of socioenvironmental exposures^22,23^. Consequently, the genetic basis of DNAm differences and the implications of genetic ancestry-specific DNAm regulation in breast cancer remain understudied.

In this study, we investigated the contribution of genetic variation to DNAm differences between breast cancer patients of African and European ancestry, focusing on the regulatory effects on gene expression. We used multi-omic data from The Cancer Genome Atlas (TCGA) breast cancer (BRCA) cohort for an integrative analysis of 578 patients of European and African ancestry. We identified differential DNAm sites and examined the regulatory effects of genetic variation on these sites. Our findings reveal additive genetic regulatory effects on ancestry-related DNAm differences, with most alleles acting as meQTLs showing differential frequencies by ancestry and acting from a long distance. Additionally, by mapping expression quantitative trait methylation (eQTMs), we found that ancestry-specific DNAm patterns form gene regulation communities affecting immune processes in breast cancer. Differential expression analysis revealed that 31% of the differentially expressed genes between African and European patients are under ancestry eQTM control. These findings highlight the significant impact of ancestry-specific genetic regulation on DNAm and its potential to confound interpretations of DNA methylation-based signatures of environmental exposures.

## Results

### Differential methylation patterns distinguishing African and European breast cancer patients are in functional parts of the genome

We performed differential DNA methylation (DNAm) analysis between women of African (AFR) and European (EUR) genetic ancestry to determine the prevalence of ancestry associated DNAm differences in the TCGA breast cancer cohort. Although there were many DNAm differences with small effect sizes, here we report results based on sites that were both significantly differentially methylated and had a large enough effect size to be considered biologically meaningful. Using these criteria, we identified 757 ancestry-associated differentially methylated sites (aDMS) between AFR and EUR women with breast cancer (Figure 2a, Figure S1a), with 31% of aDMS hypermethylated in EUR and 67% of aDMS hypermethylated in AFR women with breast cancer respectively. Analyzing the extent to which DNA methylation variation at these CpG sites is associated with genetic ancestry, we used principal components analysis (PCA) to confirm that the major axes of variation captured at these sites separated African from European patients, with PC1 and PC2 capturing 15.37% and 5.5% of the DNAm variation associated with genetic ancestry respectively (Figure S1b).

**Figure 1.**
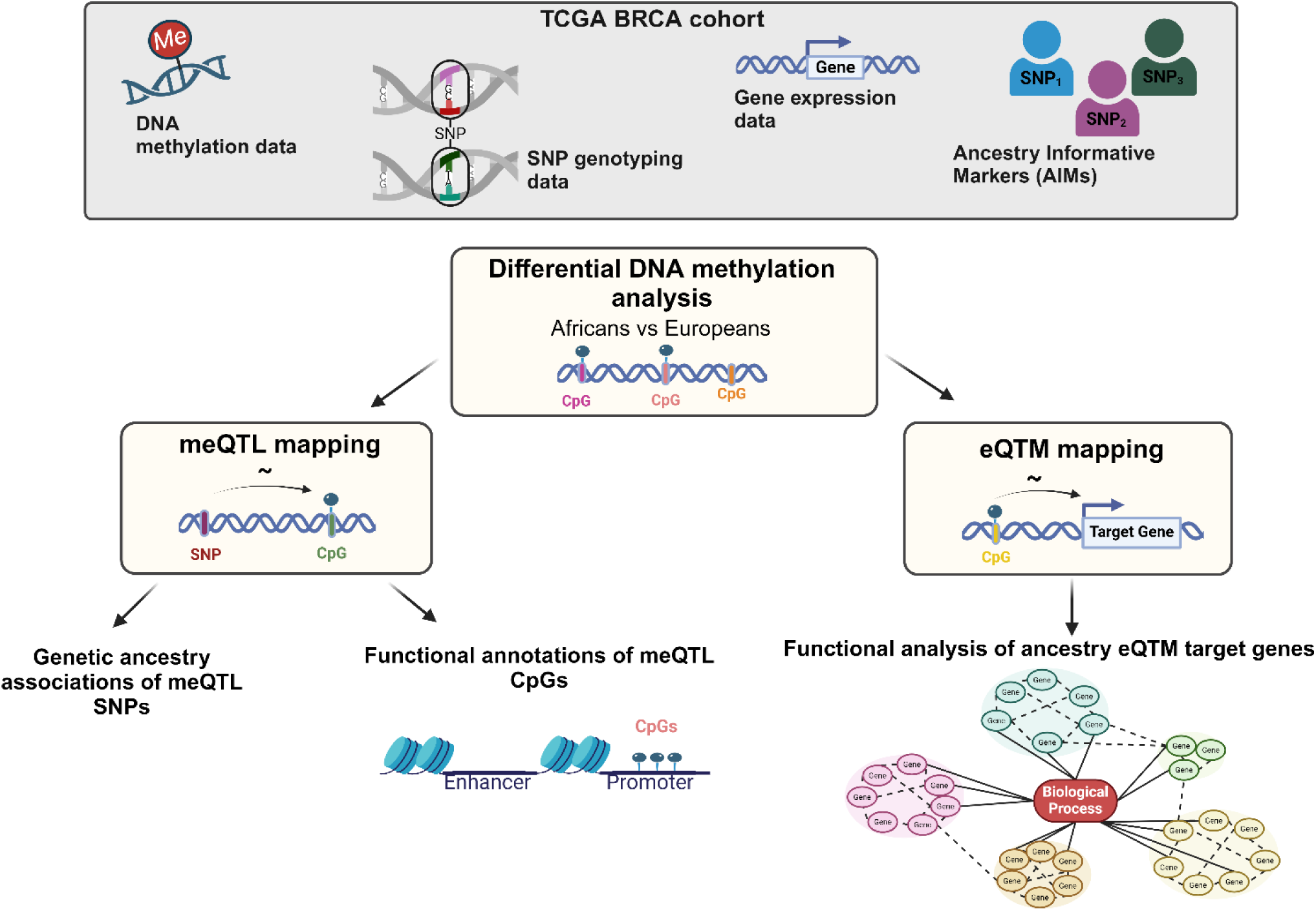
Schematic of Analysis Overview.

**Figure 2.**
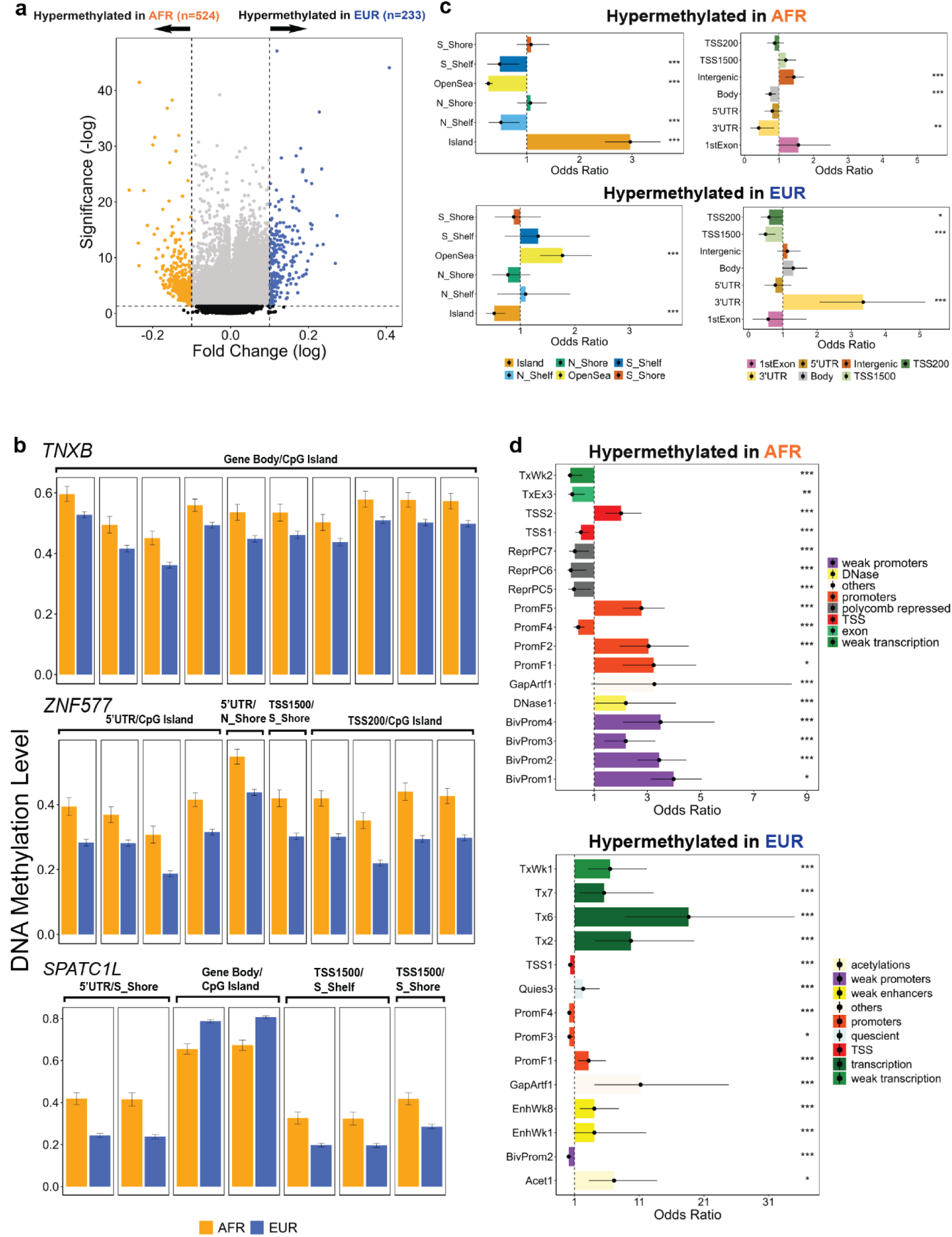
Genetic Ancestry Specific Methylation Shows Distinct Localization patterns in the TCGA BRCA Cohort. a, Differentially methylated probes between women of African (AFR) and European (EUR) ancestry with breast cancer in the TCGA cohort. Each point represents a single Illumina array probe and color-coding highlights probes that showed statistically significant differences in methylation between the two groups (blue: q-value (Benjamini-Hochberg adjusted p-value) < 0.05, effect size < 0.1; red: unadjusted p-value < 0.05 and effect size ≥ 0.1). Positive Log Fold Change values indicate hypermethylation in EUR, while negative Log Fold Change values indicate hypermethylation in AFR. **b,** Mean average beta methylation values across ancestries for the 3 genes associated with the highest number of differentially methylated probes by ancestry. **c,** Enrichment odds ratios for the 757 differentially methylated probes across CpG Island and gene structure elements examined separately for probes hypermethylated in AFR and EUR. 95% confidence are intervals also indicated. CpG annotated percentages were compared to the respective proportion of the 486,435 annotated Illumina array probes (fisher’s exact test). All associations are displayed. **d,** Enrichment odds ratios for the 757 differentially methylated probes across 100 different chromatin states examined separately for probes hypermethylated in AFR and EUR. 95% confidence intervals are also indicated. CpG annotated percentages were compared to the respective proportion of the 486,435 annotated Illumina array probes (fisher’s exact test). Only significant associations are displayed (p-value < 0.05). *For all associations, a p-value < 0.05 and* ≥ *0.03 is indicated with “*”, a p-value < 0.03 and* ≥ *0.01 is indicated with “**”, while a p-value < 0.01 is indicated with “*”*.

aDMS display differential enrichment scores across functional regions of the genome with sites hypermethylated in AFR individuals more likely to be found in intergenic regions and CpG Islands and depleted in 3’ UTRs and gene bodies (Figure 2c). Statistically significant enrichment of EUR-specific methylation is conversely observed in 3’ Untranslated regions (UTRs) and open sea regions. Additionally, EUR aDMS were depleted in CpG Islands and promoter regions (Figure 2c). aDMS localization across chromatin states showed enrichment of AFR aDMS primarily in promoters while EUR aDMS were mainly detected in genomic regions that are transcribed (Figure 2d). Additionally, 6% of aDMS are in open chromatin regions previously shown to exhibit African ancestry-specific accessibility patterns in breast cancer cell lines^24^. Interestingly, the level of differential accessibility in these regions was negatively correlated with the differential DNAm value at the aDMS (Pearson correlation coefficient -0.53, p-value 0.003) (Supplementary Table 2). This finding suggests that chromatin regions that are more accessible in one population group than in the other, are also hypomethylated, pointing to the potential for regulatory elements to bind to these regions and exert their influence on gene expression. Taken together, these results have functional implications for ancestry-associated differential methylation because of the differential localization patterns across genomic regions that play critical regulatory roles in transcription.

When mapped to annotated genes, the aDMS recapitulated previous findings and implicated genes known to be involved in cancer-associated processes. For example, *SPATC1L* with 7 aDMS nearby (Figure 2b) was previously reported to harbor ancestry-associated DNAm differences in a pan-cancer analysis of the TCGA cohort that included the TCGA-BRCA cohort under study^21^. *TNXB* and *ZNF577,* which each had 10 aDMS associated with them, had the highest number of ancestry-associated CpGs annotated to them (Figure 2b). Intriguingly, changes in *TNXB* expression have been previously associated with breast cancer progression^25^, while methylation changes in *ZNF577* have been associated with obesity-related breast cancer^26^. These findings point to a role for ancestry-specific methylation in regulating genes implicated in breast cancer pathogenesis.

### meQTL mapping reveals long-distance genetic regulation of the ancestry associated DNAm differences

To examine whether ancestry associated DNAm differences are under genetic control, we performed meQTL mapping on the aDMS to identify polymorphic loci in the genome associated with the methylation status at these sites. We examined both local (cis-meQTLs) and long distance (trans-meQTLs) associations and identified a large number of meQTLs. Most of the genetic effects on aDMS had small effect sizes. Using a stringent effect size threshold used in previous cancer meQTL studies^27^, we report 158 cis- and2,774 trans-meQTLs, with 26.2%(198/757) of aDMS participating in at least one cis-and/or trans-meQTL (Figure 3a, Supplementary Table 3). We refer to these meQTLs as high impact meQTLs, due to their substantial effect sizes. We found that a single aDMS can be regulated by multiple meQTL SNPs (meSNPs) (Figure 3a), while the highest number of meSNPs regulating the same aDMS was 504 (Figure 3a, 3d). This suggests that the genetic effects on aDMS may be additive, aligning with previous studies reporting that genetic effects on DNAm typically display small effect sizes and act additively^28^. The 2,774 trans-meQTLs were formed by associations between 2,322 SNPs and 187 aDMS, while the 158 cis-meQTLs were formed by SNP-CpG associations between 136 SNPs and 31 aDMS. Notably, 20 of these aDMS were regulated by both cis- and trans-meQTLs. We calculated enrichment odds ratios across different functional regions of the genome to pinpoint the locations of the meQTL regulated aDMS. We found significant enrichment of meCpGs in CpG Islands and intergenic regions, with significant depletion in TSS200 regions (Figure 3b). Additionally, we found an enrichment of meCpGs in promoters and histone acetylation states based on chromatin state annotations (Figure 3b). These results suggest that genetic variation affecting ancestry-associated DNAm primarily acts via long distance mechanisms and mainly influences methylation at functionally important genomic regions.

**Figure 3.**
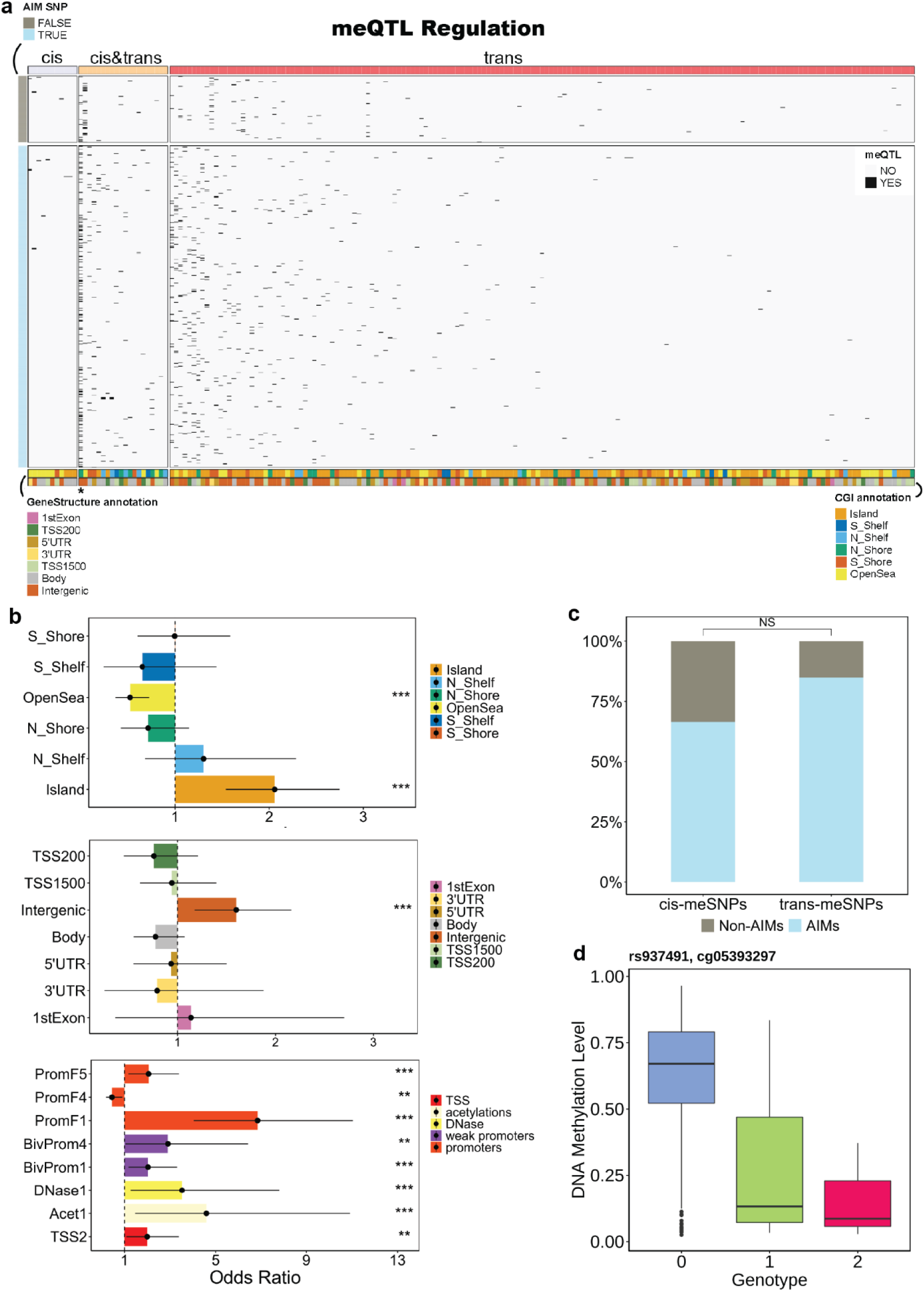
Genetic Regulation of Ancestry Associated Methylation Differences. **a,** Pleiotropic meQTL regulation of the ancestry differentially methylated sites. Number of regulatory SNPs acting as cis-and/or trans-meQTLs per ancestry differential methylated site, for the 198 CpG sites identified to participate in at least one meQTL. 11 ancestry differentially methylated sites are regulated only via cis-meQTLs, 167 are regulated by trans-meQTLs, while 20 sites are regulated by both cis- andtrans-meQTLs. The meQTL SNPs are clustered based on whether they are AIMs or not (row annotation of the heatmap). The asterisk indicates the CpG site with the highest number of meQTLs (probe cpg05393297). **b,** Enrichment odds ratios for the 132 unique meCpGs across CpG Island elements gene structure elements and 100 different chromatin states together with the 95% CI. CpG annotated percentages were compared to the respective proportion of the 486,435 annotated Illumina array probes (fisher’s exact test). Only significant associations are displayed (p-value < 0.05). **c,** Percentage of SNPs identified to be Ancestry Informative Markers across in cis- andtrans-meQTLs. **d,** meQTL plot visualizing the effect of one of the meSNPs for cg05393297 on the methylation value at this site. rs937491 has differential frequency by ancestry and it is considered an Ancestry Informative Marker (AIM). Methylation value of cg05393297 is shown for the patients with zero, one, or two copies of the allele. *For all associations, a p-value < 0.05 and* ≥ *0.03 is indicated with “*”, a p-value < 0.03 and* ≥ *0.01 is indicated with “**”, while a p-value < 0.01 is indicated with “*”*.

### The majority of meQTL SNPs regulating ancestry associated DNA methylation have differential allele frequencies by ancestry

To investigate further whether meQTL regulation of the aDMS is driven by population specific genetic variation, we examined the association between meQTL SNPs and germline genetic differences between population groups. Here we used genetic polymorphisms with varying genotype frequencies associated with genetic ancestry identified by Carrot-Zhang et al.^21^ as ancestry informative markers (AIMs). We found that 64% of the cis-meQTL SNPs and 84% of the trans-meQTL SNPs are AIMs^21^, with these differences in genotype frequencies between populations plausibly accounting for differences in methylation levels between these groups (Figure 3c). In parallel, we performed a control analysis where we ran meQTL analysis on a random set of 757 CpG sites, with no ancestry differential methylation. Here we identified no cis-meQTLs and 128 trans-meQTLs with ∼9.4% (12/128) AIM trans-meSNPs. Although the difference in AIM prevalence between cis- and trans-meQTLs was not statistically significant for the aDMS, the higher number of identified trans-meQTLs and the higher prevalence of AIM SNPs exerting their influence on methylation in trans suggests that the effects of genetic ancestry on DNAm primarily operate through long distance associations. These findings point to a role for population-specific genetic variation in regulating DNA methylation at sites that show ancestry-associated methylation patterns.

### DNAm differences by ancestry form communities of gene regulation primarily affecting immune response processes

We then sought to identify local genes that are influenced by the methylation status at aDMS to ascertain the functional impact of these ancestry-associated differences in methylation. We mapped expression quantitative trait methylation loci (eQTMs), by associating methylation at aDMS with local gene expression. 95% of the aDMS act as ancestry eQTMs, significantly influencing expression in at least one gene (eQTM target gene) in the local neighborhood (±1.5Mb window). In total, 5,097 genes were regulated by ancestry eQTMs, with additive effects observed—a single gene could be regulated by multiple ancestry eQTMs. Interestingly, as the number of eQTMs regulating a gene increased, the difference in that gene’s expression level between African and European breast cancer patients also increased (Figure 4a). Additionally, 37% of the eQTM target genes were regulated by an ancestry eQTM under the influence of at least one high-impact meQTL (meQTL-eQTM regulation), while 72% of the eQTM target genes were associated with an eQTM regulated by at least one low- or high-impact meQTL (Methods).

**Figure 4.**
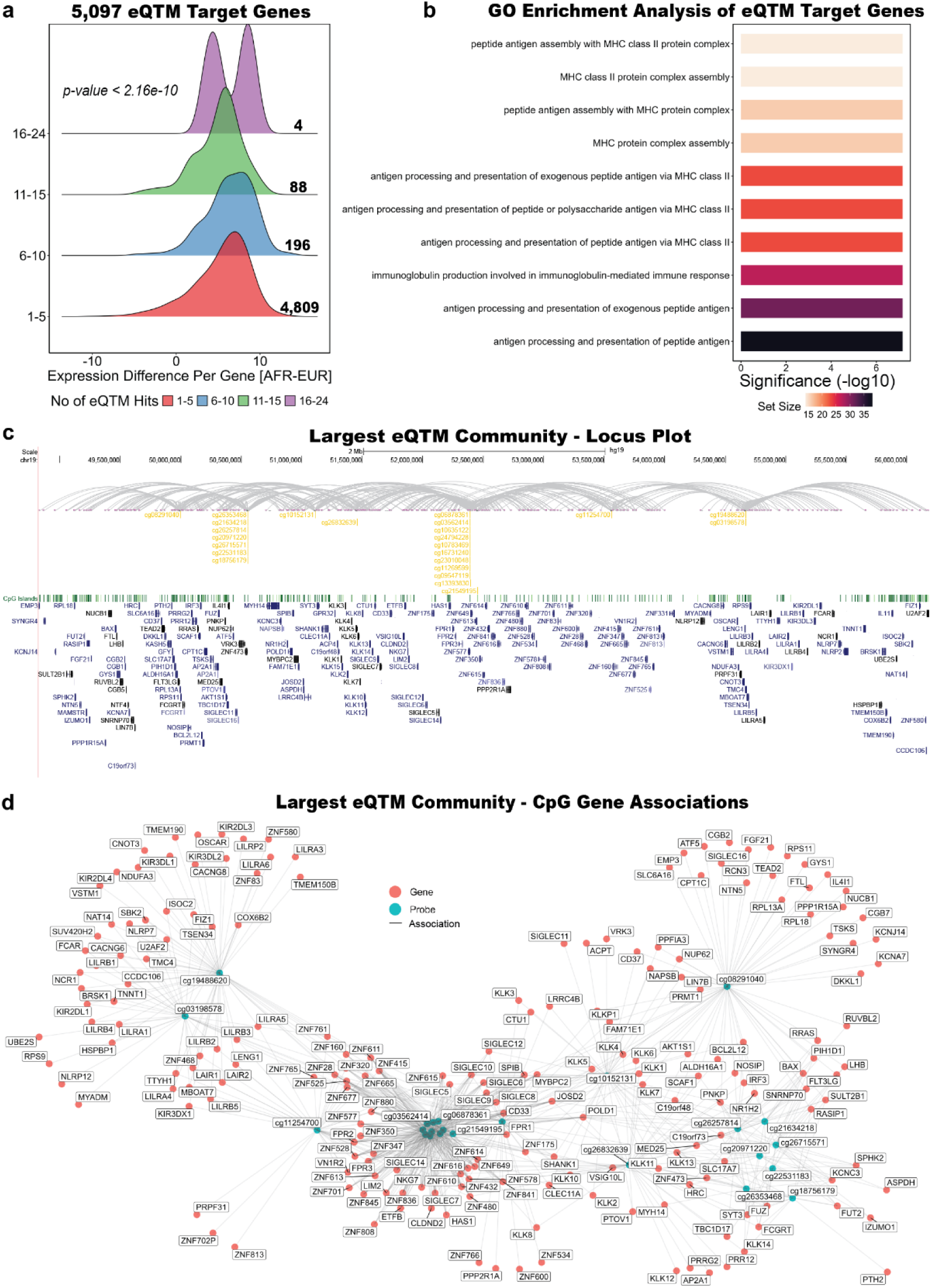
Pleiotropic and Additive Effects of Ancestry Specific Methylation on Gene Expression. **a,** Difference in gene expression between AFR and EUR increases as the number of ancestry eQTMs regulating a gene increases. Y axis represents number of regulatory eQTMs per gene, x axis represents the log2-tranformed absolute difference in gene expression between AFR and EUR. The results are visualized in groups (genes regulated by 1 to 5, 6 to 10, 11 to 15, and 16 to 24 eQTMs). Number of genes in each group is indicated on the right of the distribution curve. **b**, First ten most significantly eQTM enriched Biological Process Gene Ontologies. Bar plot is ordered based on decreasing significance (-log10 adjusted p-value). The set size of each enriched GO is also indicated. **c,** Locus plots of the largest eQTM community. Representation of the ancestry eQTMs and their target genes in the HLA locus. Ancestry eQTMs are highlighted in yellow, and gray arrows indicate the genes regulated by each ancestry eQTM. Pink dots represent the position of the eQTM target genes, while gene names are highlighted in a color scale from light blue to black (**light blue**: non-RefSeq transcripts, **medium blue**: other RefSeq transcripts, **dark blue**: transcript has been reviewed or validated by either the RefSeq, SwissProt or CCDS staff, **black**: feature has a corresponding entry in the Protein Data Bank (PDB)). CpG Islands in the locus and their respective positions are also indicated. **d,** Network graph of the largest eQTM community. The ancestry eQTMs are highlighted in blue, while the eQTM target genes in the community are highlighted in orange.

We considered all the identified eQTMs and their target genes as an extensive eQTM network. Analysis of this network revealed that eQTMs close to each other in the genome form regulatory communities, influencing expression of genes with similar functions in that same locus. We identified 278 eQTM communities, the largest of which had 24 ancestry eQTMs regulating 203 genes within a ∼7.4Mb window on chromosome 19 (Figure 4c, 4d). Unsurprisingly, the eQTM targets in this community were primarily genes involved in transcriptional regulation and RNA/peptide biosynthesis (Figure S2).

The second largest eQTM community included the HLA locus on chromosome 6, where 23 ancestry eQTMs regulated 173 eQTM target genes, including multiple HLA genes (Figure S3a). Clustering the eQTM target genes at the HLA locus based on their correlation with ancestry eQTMs identified three main clusters. Interestingly, genetic and transcriptional variation in HLA genes, specifically associated with African ancestry, has been extensively reported previously^29–31^. Most of the HLA genes were in cluster 3, showing a consistent expression-methylation correlation pattern across the ancestry eQTMs in that community (Figure S3b).

The biological processes most enriched across all 5,097 ancestry eQTM target genes were those associated with the regulation of immune responses (Figure 4b, Figure S4a). The most significantly enriched processes were associated with MHC class II protein complex assembly, driven primarily by the HLA genes (Figure S4b). Although the HLA locus eQTM community ranked second in size, it had the highest number of eQTM associations, indicating high pleiotropy, with each ancestry eQTM regulating many genes within the network.

### Differential expression of eQTM target genes

To understand whether ancestry-associated DNAm differences affect gene expression and consequently influence breast cancer phenotypes, we next quantified the proportion of eQTM target genes that are differentially expressed between Africans and Europeans. We identified 374 differentially expressed genes between the two population groups; 116 of these genes were associated with at least one ancestry eQTM. Notably, we observed that for 73.3% of the differentially expressed eQTM target genes, the regulating eQTM was influenced by at least one high- or low-impact meQTL (meQTL-eQTM regulation) (Supplementary Table 5).

Although only 2.3% (116/5,097) of the eQTM target genes exhibited differential expression between AFR and EUR breast cancer patients, 31% (116/374) of the differentially expressed genes by ancestry were regulated by ancestry-specific eQTMs. Additionally, 23% of all differentially expressed genes were under the combined influence of ancestry-specific meQTL and eQTMs. This finding highlights the non-negligible regulatory role of ancestry-specific genetic and epigenetic variation in gene expression differences in disease contexts such as breast cancer.

Our findings are in line with previous results. Of the 59 genes identified by Carrot-Zhang et al.^21^ as African ancestry-associated differentially expressed genes in breast cancer, 53 were also detected as differentially expressed in our study. Interestingly, 20% (11/53) of these genes were under ancestry eQTM control, while almost all of them (9 of the 11) were under joint meQTL-eQTM regulation (Supplementary Table 5). This reinforces the importance of considering ancestry-specific genetic regulation of DNAm in understanding gene expression differences and their implications in breast cancer.

## Discussion

Population-level DNA methylation (DNAm) differences in breast cancer have primarily been used in molecular subclassification studies or determining the molecular effects of an adverse lifestyle or exposure to adverse environments. However, the contribution of genetic variation to DNA methylation variability in these contexts has yet to be fully explored. In this study, we show clearly that DNAm differences between individuals of African and European ancestry in the TCGA BRCA cohort are strongly associated with genetic variation.

Ancestry-specific methylation exhibits distinct localization patterns across the genome. African-specific hypermethylation is mainly observed in promoters and CpG islands, while European-specific hypermethylated sites are detected within genes, transcribed regions, and 3’ UTRs. Ancestry differentially methylated sites often form regulatory gene communities, acting as local eQTMs and regulating genes with similar functions within the same locus. Immune regulatory processes, especially MHC class II and antigen processing, are most affected by ancestry eQTMs, with HLA genes driving the enrichment in these specific biological processes (Figure S4b). Approximately 72% of eQTM-associated DMS are regulated by highly pleiotropic meQTLs, dominated by trans-meQTLs with meSNPs with differential frequencies by ancestry. More importantly, genetic ancestry-specific DNAm regulation is linked to gene expression changes, with nearly one-third of differentially expressed genes between African and European breast cancer patients under joint ancestry meQTL-eQTM control.

Our findings indicate that population-level genomic differences are strongly linked to distinct DNAm patterns in breast cancer patients. Consequently, studies that consider DNAm solely as a measure of adverse environmental exposures in cancer, without accounting for population stratification, risk confounding. Interestingly, the genes most significantly associated with CpG sites under genetic control are involved in transcriptional regulation broadly and the regulation of immune responses. HLA genes, for which differences specifically associated with African ancestry have been widely reported^29–31^, are the most enriched for genetic ancestry-related regulatory effects. Additionally, we found that the majority of meQTLs regulating aDMS are SNPs with differential frequencies by ancestry. Therefore, filtering for rare variants (MAF < 0.01) in meQTL studies with individuals from diverse ancestry backgrounds is an analytic step that can potentially lead to biased results, as variants rare in European ancestry-enriched cohorts are not necessarily rare across all population groups.

Our study has several limitations that may limit our ability to adequately determine the role of ancestry-specific effects on DNA methylation. The primary limitation is statistical power, with 487 tumor samples from individuals of European ancestry and 91 from individuals of African ancestry, potentially affecting differential methylation and meQTL analysis. Additionally, the study was limited to CpG sites captured on the Illumina Human Methylation 450k array, which mainly covers CpGs in gene promoters and CpG islands, with limited coverage of other regulatory regions, such as those in non-coding parts of the genome^32^. Previous studies have reported that CpGs associated with DNAm patterns between populations are primarily located in non-coding genomic regions^17,33^. Consistent with these findings, we identified significant African-specific hypermethylation in intergenic regions and European-specific hypermethylation in open sea regions (Figure 2c). Deeper base resolution coverage of CpGs genome-wide would likely have increased the number of identified ancestry-associated methylation differences and their regulatory architecture (meQTLs, and eQTMs). A critical limitation is the availability of additional public multi-omic breast cancer datasets that jointly characterize the genome, methylome and transcriptome in individuals of diverse genetic ancestry to further validate the results reported here.

Furthermore, we were not able to investigate the potential interactions between genetic variation and environmental exposures in breast cancer as TCGA does not have exposome information annotated on that cohort. Future research studies will need to jointly collect both the requisite biospecimens and exposure metadata to enable the study of gene-by-environment interactions in breast cancer. This will allow us to fully understand the relative contributions of population-level genetic differences on DNAm variability in breast cancer and its functional impact in cancer. Additionally, local ancestry effects should be considered to develop a comprehensive picture of these interactions.

Taken together, our findings provide new insights into how population level genetic variation can influence epigenetic changes that can affect immune regulation in breast cancer. These results enhance our understanding of the basis of ancestry-associated DNAm differences in breast cancer while also suggesting a link between genetic ancestry-regulated DNAm and the more aggressive and inflammatory breast cancer phenotypes observed in individuals of African descent.

**Figure S1.**
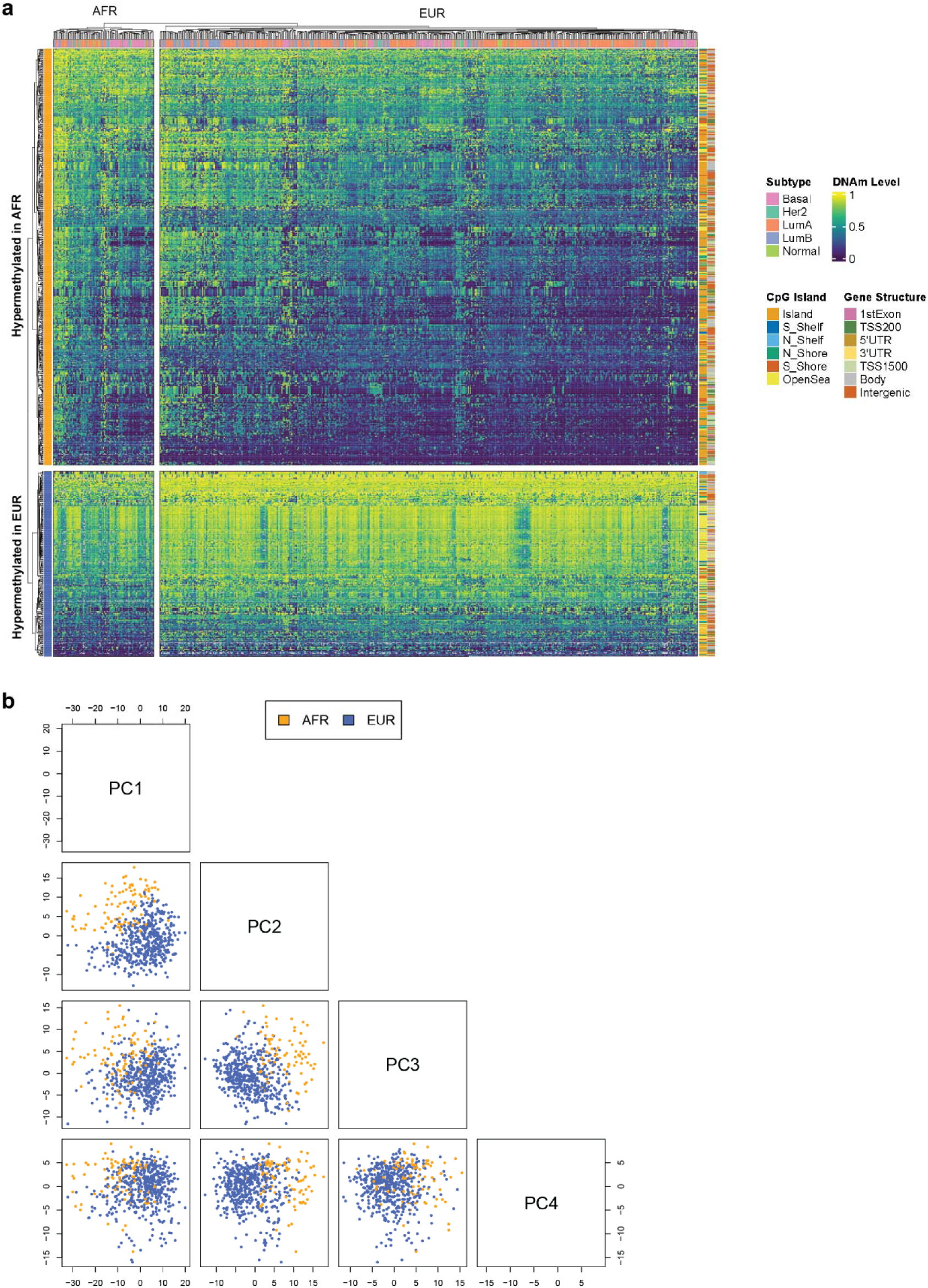
Distinct DNAm Patterns Between Breast Cancer Patients of African (AFR) and European (EUR) Ancestry in the TCGA Cohort. **a,** Differentially methylated sites (DMS) between women of AFR and EUR ancestry with breast cancer in the TCGA cohort. Each point represents a single Illumina array probe, and color-coding highlights probes that showed statistically significant differences in methylation between the two groups (blue: q-value (Benjamini-Hochberg adjusted p-value) < 0.05, effect size < 0.1; red: unadjusted p-value < 0.05 and effect size ≥ 0.1). Positive Log Fold Change values indicate hypermethylation in EUR, while negative Log Fold Change values indicate hypermethylation in AFR. **b,** Principal Component Analysis (PCA) biplots on the beta methylation values of the 757 differentially methylated probes for AFR (orange) and EUR (blue) breast cancer patients in the cohort.

**Figure S2.**
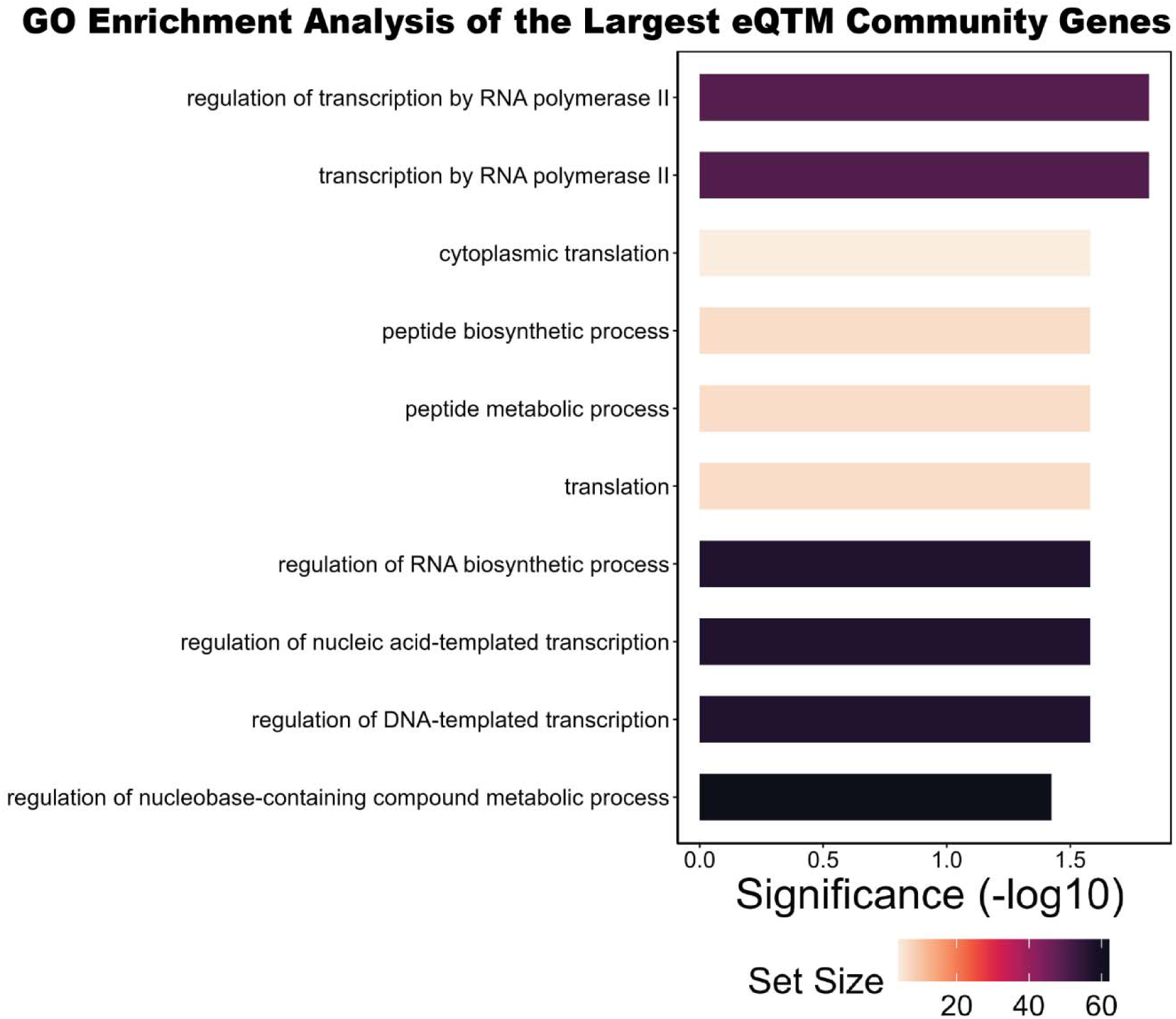
Target Genes in the Largest eQTM Community are Involved in Essential Molecular Processes. Enriched Biological Process Gene Ontologies among the target genes of the largest eQTM community.

**Figure S3.**
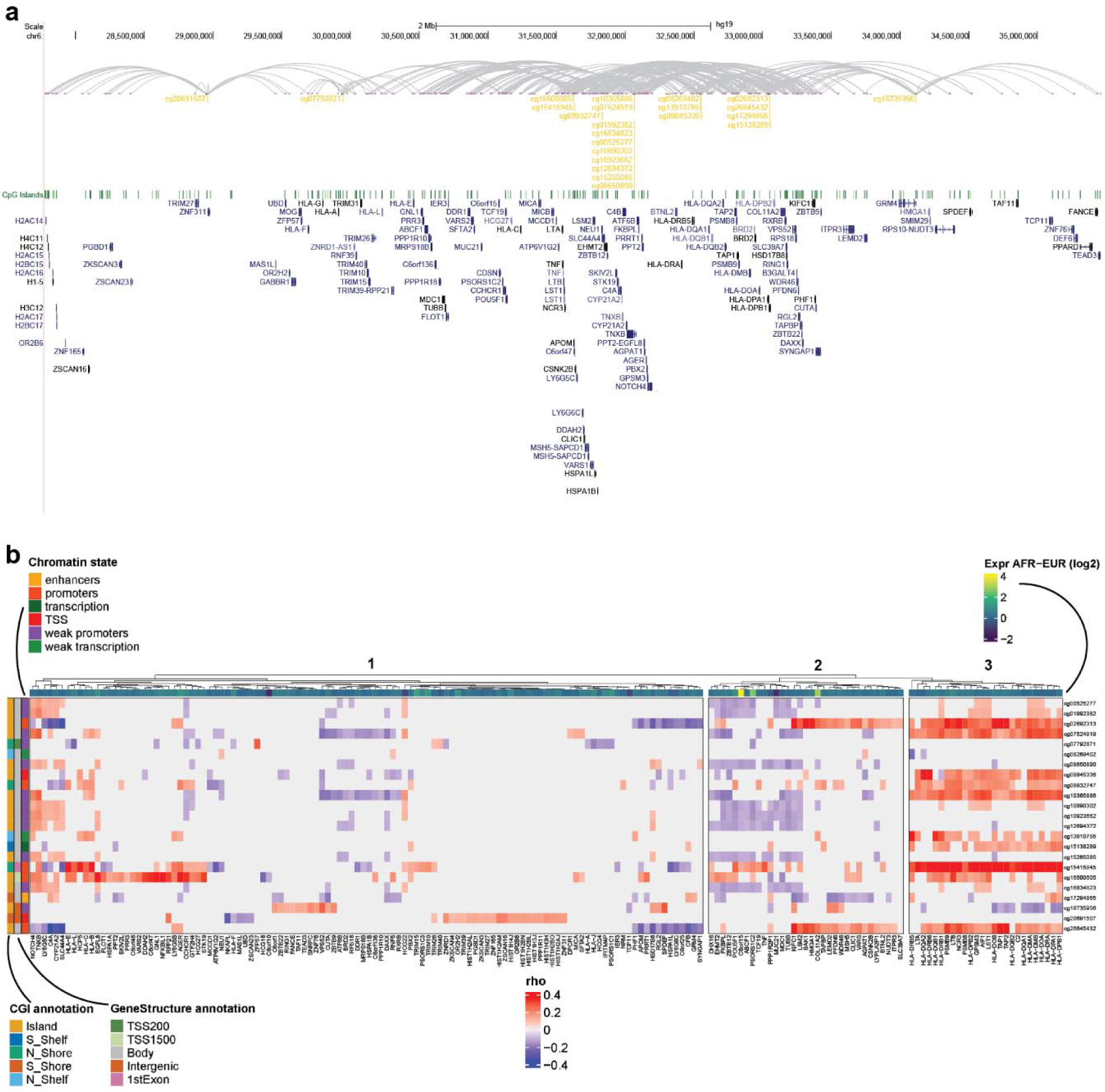
Chromosome 6 HLA Locus Is under Ancestry Specific Methylation Control. **a,** Second largest eQTM community: Chromosome 6 HLA locus. Representation of the ancestry eQTMs and their target genes in the HLA locus. Ancestry eQTMs are highlighted in yellow, and gray arrows indicate the genes regulated by each ancestry eQTM. Pink dots represent the position of the eQTM target genes, while gene names are highlighted in a color scale from light blue to black (**light blue**: non-RefSeq transcripts, **medium blue**: other RefSeq transcripts, **dark blue**: transcript has been reviewed or validated by either the RefSeq, SwissProt or CCDS staff, **black**: feature has a corresponding entry in the Protein Data Bank (PDB)). CpG Islands in the locus and their respective positions are also indicated. **b,** Methylation ∼ Expression Correlation Heatmap for the second largest eQTM community. CpG sites that act as eQTMs in the chromosome 6 HLA locus eQTM community are shown on the right. Spearman correlation coefficient (rho) between methylation value in the eQTMs and expression values across all genes in the community is visualized on a scale from blue (negative correlation) to red (positive correlation). CpG Island (CGI), Gene Structure, and Chromatin State annotations are also provided for the eQTMs (row annotations). The difference of log2 transformed expression values of AFR and EUR patients is also displayed for every gene in the community (column annotations).

**Figure S4.**
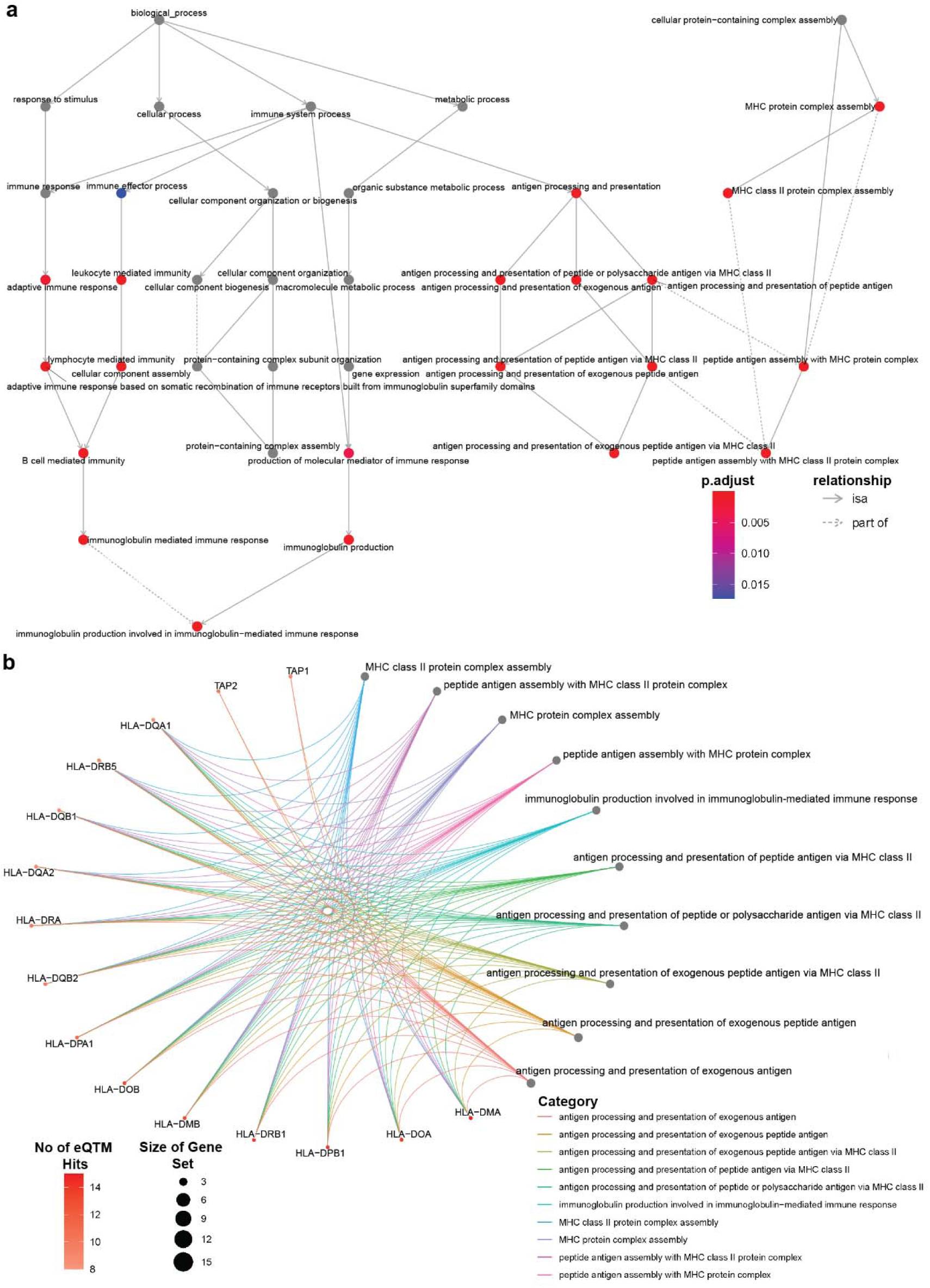
Ancestry eQTMs Primarily Influence Regulatory Immune Response Processes. **a,** Gene Set Enrichment Analysis on the eQTM target genes. eQTM enriched Biological Processes (BP) presented in a hierarchical layout. The terms statistically enriched in our analysis are colored in a scale from blue to red (based on their p-value), while gray dots are visualizing the structure of the Ontology. The arrowhead lines represent the direction of the relationship (*is a*: parent to child node), while the dotted edges (*part of*) represent inferred relationships between the Gene Ontology terms. **b,** Circular Enrichment plot for the 10 most significantly ancestry eQTM enriched BP Ontologies. The specific genes significantly enriched for eQTM regulation in each GO category are shown on the left, while colored curves connect the genes with the Ontologies they belong to.

## Methods

### DNA Methylation Data

We utilized the preprocessed TCGA Pan-cancer methylation matrix (Illumina Human Methylation 450k array data) from Carrot-Zhang et al.^21^ Using the sample-TCGA cancer type mapping information derived from Sanchez-Vega et al.^34^, we selected breast cancer (BRCA) samples from the pan-cancer matrix (873 samples). Sample quality information such as age was derived from the PanCanAtlas Publications Genomic Data Commons archive^34^. We excluded samples with unknown tumor purity or age at initial pathological diagnosis from further analysis. Our final cohort consisted of 617 BRCA samples.

### SNP Genotyping Data

We used TCGA SNP genotyping data based on preprocessing and quality control steps described in Sayaman et al.^35^. meQTL analyses were based on the stranded 1000 Genome Project (1000G) imputed genotyping calls filtered for BRCA samples that also had DNAm data available (n=554).

### Gene Expression Data

In this study, we used the preprocessed and batch-corrected TCGA Pan-cancer mRNA normalized data (https://gdc.cancer.gov/about-data/publications/pancanatlas). Using the tumor type and subtype information from Sanchez-Vega et al.^34^, we subset this matrix to select TCGA breast cancer samples for this study.

### Global Genetic Ancestry Assignment

We used the ancestry assignment for each sample in the BRCA cohort as previously defined by Carrot-Zhang et al.^21^. This method is based on consensus ancestry calls from multiple approaches including calls made by ADMIXTURE^36^. For our analyses we retained samples with admixture coefficient <20% for African or European ancestry.

### Differential DNA Methylation Analysis

We performed differential DNAm analysis using a multivariate linear regression model (*limma* R package^37^), that modeled the beta value of each Illumina 450K probe (DNA methylation level at a given CpG site) as the outcome variable. Explanatory variables included genetic ancestry group (African and European ancestry (578 samples)), age, tumor purity, and BRCA subtype. We considered probes with Benjamini–Hochberg^38^ adjusted p-value < 0.05 and effect size ≥ 0.1 as significantly differential methylated. We excluded 47 “rs” probes, which are used as positive controls, as their measurements are directly influenced by SNPs (‘‘SNP masked’’).

Enrichment odds ratios for the differentially methylated probes across different genomic regions were computed against the 486,435 Illumina Human Methylation 450k array probes using Fisher’s exact test. For probes with multiple annotations, we selected one according to the following priority order: TSS200 > TSS1500 > 5′UTR > 1st Exon > Body > 3′ UTR > Intergenic^39^. We followed the same strategy to estimate enrichment odds ratios for the ancestry meQTL CpGs.

To investigate enrichment of ancestry CpGs across different chromatin states, we used the full-stack chromatin state annotation of the genome generated by Vu et al.^40^ Using these universal chromatin state annotations, we annotated ancestry probes to chromatin states and compared them to the respective annotations we generated for all Illumina array probes using Fisher’s exact test.

### Ancestry Associated Differentially Methylated Sites in Differentially Accessible Chromatin Regions

We investigated the differential accessible chromatin regions between African and European triple negative breast cancer cell lines identified by Harris et al.^24^, examining potential overlap with the ancestry DMS we detected. As differentially accessible regions by ancestry we considered those with FDR ≤ 0.05 and signal fold change ≥ 2. Since the differential chromatin accessibility results were reported based on GRch38 genome version, we used the LiftOver UCSC Genome Browser tool^41^ to derive the respective positions on GRch37 genome version, as all analyses generated in our study were based on GRch37.

### meQTL Analysis

We performed meQTL analysis on 747 of the 757 differentially methylated probes, excluding the 10 probes located at sex chromosomes. We used the tensorQTL software^42^ for meQTL identification applying the analysis to 554 TCGA BRCA samples with both SNP genotyping and methylation data available. In the meQTL model, we controlled for the first three PEER factors^43^ (quantile-normalized DNAm residuals were obtained) as well as age, tumor purity, and cancer subtype. The cis-meQTL window was 1Mb. We considered meQTLs with p-value < 2e-5 and effect size ≥ 0.25 as high impact ancestry-associated meQTLs. The effect size threshold was selected based on previously published cancer meQTL studies^27^, as well as results of sensitivity analysis we carried out where we examined how the number of meQTLs changes as the effect size varies (data not shown). Low impact meQTLs were identified based on effect size threshold of ≥ 0.1, a cutoff widely used in different types of omics analyses.

Probe annotations for the meQTL CpGs were derived from the Illumina Human Methylation 450k array annotation file (https://webdata.illumina.com/downloads/productfiles/humanmethylation450/humanmethylation450_15 <u>017482_v1-2.csv</u>). Odds ratios for CpG localization were computed based on comparison with the 486,435 annotated Illumina array probes.

For the control meQTL analysis we used the same breast cancer samples and ran the analysis on a randomly selected set of Illumina array probes, which didn’t show any significant differentially methylation patterns by ancestry (adjusted p-value > 0.05).

### eQTM Analysis

For association between DNAm and proximal gene expression we chose a ±1.5Mb window, considering the CpG site as the center of the window^44^. Using samples with gene expression and methylation data available, we calculated Spearman correlation between the beta value of each of the differential methylated probes and the expression levels of proximal genes (genes within a ±1.5Mb window)^44^. We identified CpGs acting as eQTMs at Benjamini–Hochberg^38^ adjusted p-value < 0.05.

To identify communities of regulatory CpGs and target genes across the identified eQTMs, we used the “NetworkX” python package^45^. eQTM networks were identified using a greedy modularity maximization algorithm, which uses Clauset-Newman-Moore greedy modularity maximization to find the community partition with the largest modularity^46^.

### Differential Expression Analysis

We log2-transformed transformed the gene expression data and filtered out genes not expressed in at least 20% of the samples. We performed multivariate regression analysis (*limma* R package^37^) comparing African and European samples (691 samples) and modeling covariables for breast cancer subtype and sequencing/processing batch^21^. We obtained batch information from the plate ID portion of the TCGA barcode. We applied a threshold of Benjamini–Hochberg^38^ adjusted p-value (q-value) < 0.05 and effect size ≥ 0.1 to identify statistically significant differentially expressed genes.

### Gene Set Enrichment Analysis of eQTM target genes

We performed Gene Ontology (GO)^47^ analysis to explore the biological functions of the genes affected by ancestry associated eQTMs. We used the “GSEA” function of the *clusterProfiler* R package^48^. To identify significantly enriched GOs we chose a threshold of Benjamini-Hochberg^38^ adjusted p-value < 0.05 and 1 for the weight of each step in the Biological Processes GO hierarchy.

## References

1. Sung H, Ferlay J, Siegel RL, et al. Global Cancer Statistics 2020: GLOBOCAN Estimates of Incidence and Mortality Worldwide for 36 Cancers in 185 Countries. CA Cancer J Clin. 2021;71(3):209-249. doi:10.3322/caac.21660

2. Siddharth S, Sharma D. Racial Disparity and Triple-Negative Breast Cancer in African-American Women: A Multifaceted Affair between Obesity, Biology, and Socioeconomic Determinants. Cancers. 2018;10(12):514. doi:10.3390/cancers10120514

3. Giaquinto AN, Sung H, Miller KD, et al. Breast Cancer Statistics, 2022. CA Cancer J Clin. 2022;72(6):524-541. doi:10.3322/caac.21754

4. Martini R, Delpe P, Chu TR, et al. African Ancestry–Associated Gene Expression Profiles in Triple-Negative Breast Cancer Underlie Altered Tumor Biology and Clinical Outcome in Women of African Descent. Cancer Discov. 2022;12(11):2530–2551. doi:10.1158/2159-8290.CD-22-0138

5. Minas TZ, Kiely M, Ajao A, Ambs S. An overview of cancer health disparities: new approaches and insights and why they matter. Carcinogenesis. 2021;42(1):2–13. doi:10.1093/carcin/bgaa121

6. Zavala VA, Bracci PM, Carethers JM, et al. Cancer health disparities in racial/ethnic minorities in the United States. Br J Cancer. 2021;124(2):315–332. doi:10.1038/s41416-020-01038-6

7. Wang S, Pitt JJ, Zheng Y, et al. Germline variants and somatic mutation signatures of breast cancer across populations of African and European ancestry in the US and Nigeria. Int J Cancer. 2019;145(12):3321–3333. doi:10.1002/ijc.32498

8. Polak P, Kim J, Braunstein LZ, et al. A mutational signature reveals alterations underlying deficient homologous recombination repair in breast cancer. Nat Genet. 2017;49(10):1476–1486. doi:10.1038/ng.3934

9. Dietze EC, Sistrunk C, Miranda-Carboni G, O’Regan R, Seewaldt VL. Triple-negative breast cancer in African-American women: disparities versus biology. Nat Rev Cancer. 2015;15(4):248–254. doi:10.1038/nrc3896

10. Jiagge E, Jin DX, Newberg JY, et al. Tumor sequencing of African ancestry reveals differences in clinically relevant alterations across common cancers. Cancer Cell. 2023;41(11):1963–1971.e3. doi:10.1016/j.ccell.2023.10.003

11. Suelves M, Carrió E, Núñez-Álvarez Y, Peinado MA. DNA methylation dynamics in cellular commitment and differentiation. Brief Funct Genomics. Published online June 8, 2016:elw017. doi:10.1093/bfgp/elw017

12. Cavalli G, Heard E. Advances in epigenetics link genetics to the environment and disease. Nature. 2019;571(7766):489-499. doi:10.1038/s41586-019-1411-0

13. Smith AK, Kilaru V, Kocak M, et al. Methylation quantitative trait loci (meQTLs) are consistently detected across ancestry, developmental stage, and tissue type. BMC Genomics. 2014;15(1):145. doi:10.1186/1471-2164-15-145

14. Carja O, MacIsaac JL, Mah SM, et al. Worldwide patterns of human epigenetic variation. Nat Ecol Evol. 2017;1(10):1577–1583. doi:10.1038/s41559-017-0299-z

15. Heyn H, Moran S, Hernando-Herraez I, et al. DNA methylation contributes to natural human variation. Genome Res. 2013;23(9):1363–1372. doi:10.1101/gr.154187.112

16. Hawe JS, Wilson R, Schmid KT, et al. Genetic variation influencing DNA methylation provides insights into molecular mechanisms regulating genomic function. Nat Genet. 2022;54(1):18–29. doi:10.1038/s41588-021-00969-x

17. Husquin LT, Rotival M, Fagny M, et al. Exploring the genetic basis of human population differences in DNA methylation and their causal impact on immune gene regulation. Genome Biol. 2018;19(1):222. doi:10.1186/s13059-018-1601-3

18. Nishiyama A, Nakanishi M. Navigating the DNA methylation landscape of cancer. Trends Genet. 2021;37(11):1012–1027. doi:10.1016/j.tig.2021.05.002

19. Batra RN, Lifshitz A, Vidakovic AT, et al. DNA methylation landscapes of 1538 breast cancers reveal a replication-linked clock, epigenomic instability and cis-regulation. Nat Commun. 2021;12(1):5406. doi:10.1038/s41467-021-25661-w

20. de Almeida BP, Apolónio JD, Binnie A, Castelo-Branco P. Roadmap of DNA methylation in breast cancer identifies novel prognostic biomarkers. BMC Cancer. 2019;19(1):219. doi:10.1186/s12885-019-5403-0

21. Carrot-Zhang J, Chambwe N, Damrauer JS, et al. Comprehensive Analysis of Genetic Ancestry and Its Molecular Correlates in Cancer. Cancer Cell. 2020;37(5):639–654.e6. doi:10.1016/j.ccell.2020.04.012

22. Ahmad A, Azim S, Zubair H, et al. Epigenetic basis of cancer health disparities: Looking beyond genetic differences. Biochim Biophys Acta BBA -Rev Cancer. 2017;1868(1):16–28. doi:10.1016/j.bbcan.2017.01.001

23. Jenkins BD, Rossi E, Pichardo C, et al. Neighborhood Deprivation and DNA Methylation and Expression of Cancer Genes in Breast Tumors. JAMA Netw Open. 2023;6(11):e2341651. doi:10.1001/jamanetworkopen.2023.41651

24. Harris AR, Panigrahi G, Liu H, et al. Chromatin Accessibility Landscape of Human Triple-negative Breast Cancer Cell Lines Reveals Variation by Patient Donor Ancestry. Cancer Res Commun. 2023;3(10):2014–2029. doi:10.1158/2767-9764.CRC-23-0236

25. Liot S, Aubert A, Hervieu V, et al. Loss of Tenascin-X expression during tumor progression: A new pan-cancer marker. Matrix Biol Plus. 2020;6–7:100021. doi:10.1016/j.mbplus.2020.100021

26. Crujeiras AB, Diaz-Lagares A, Stefansson OA, et al. Obesity and menopause modify the epigenomic profile of breast cancer. Endocr Relat Cancer. Published online July 2017:351–363. doi:10.1530/ERC-16-0565

27. Gong J, Wan H, Mei S, et al. Pancan-meQTL: a database to systematically evaluate the effects of genetic variants on methylation in human cancer. Nucleic Acids Res. 2019;47(D1):D1066–D1072. doi:10.1093/nar/gky814

28. DNA methylation provides molecular links underlying complex traits. Nat Genet. 2023;55(1):12-13. doi:10.1038/s41588-022-01249-y

29. Mentzer AJ, Dilthey AT, Pollard M, et al. High-resolution African HLA resource uncovers HLA-DRB1 expression effects underlying vaccine response. Nat Med. 2024;30(5):1384–1394. doi:10.1038/s41591-024-02944-5

30. Vince N, Limou S, Daya M, et al. Association of HLA-DRB1∗09:01 with tIgE levels among African-ancestry individuals with asthma. J Allergy Clin Immunol. 2020;146(1):147–155. doi:10.1016/j.jaci.2020.01.011

31. Tshabalala M, Mellet J, Vather K, et al. High Resolution HLA ∼A, ∼B, ∼C, ∼DRB1, ∼DQA1, and ∼DQB1 Diversity in South African Populations. Front Genet. 2022;13:711944. doi:10.3389/fgene.2022.711944

32. Infinium HumanMethylation450 BeadChip.

33. Li B, Aouizerat BE, Cheng Y, et al. Incorporating local ancestry improves identification of ancestry-associated methylation signatures and meQTLs in African Americans. Commun Biol. 2022;5(1):401. doi:10.1038/s42003-022-03353-5

34. Sanchez-Vega F, Mina M, Armenia J, et al. Oncogenic Signaling Pathways in The Cancer Genome Atlas. Cell. 2018;173(2):321–337.e10. doi:10.1016/j.cell.2018.03.035

35. Sayaman RW, Saad M, Thorsson V, et al. Germline genetic contribution to the immune landscape of cancer. Immunity. 2021;54(2):367–386.e8. doi:10.1016/j.immuni.2021.01.011

36. Alexander DH, Novembre J, Lange K. Fast model-based estimation of ancestry in unrelated individuals. Genome Res. 2009;19(9):1655–1664. doi:10.1101/gr.094052.109

37. Ritchie ME, Phipson B, Wu D, et al. limma powers differential expression analyses for RNA-sequencing and microarray studies. Nucleic Acids Res. 2015;43(7):e47–e47. doi:10.1093/nar/gkv007

38. Benjamini Y, Hochberg Y. Controlling the False Discovery Rate: A Practical and Powerful Approach to Multiple Testing. J R Stat Soc Ser B Methodol. 1995;57(1):289–300.

39. Shang L, Zhao W, Wang YZ, et al. meQTL mapping in the GENOA study reveals genetic determinants of DNA methylation in African Americans. Nat Commun. 2023;14(1):2711. doi:10.1038/s41467-023-37961-4

40. Vu H, Ernst J. Universal annotation of the human genome through integration of over a thousand epigenomic datasets. Genome Biol. 2022;23(1):9. doi:10.1186/s13059-021-02572-z

41. Hinrichs AS. The UCSC Genome Browser Database: update 2006. Nucleic Acids Res. 2006;34(90001):D590–D598. doi:10.1093/nar/gkj144

42. Taylor-Weiner A, Aguet F, Haradhvala NJ, et al. Scaling computational genomics to millions of individuals with GPUs. Genome Biol. 2019;20(1):228. doi:10.1186/s13059-019-1836-7

43. Stegle O, Parts L, Piipari M, Winn J, Durbin R. Using probabilistic estimation of expression residuals (PEER) to obtain increased power and interpretability of gene expression analyses. Nat Protoc. 2012;7(3):500–507. doi:10.1038/nprot.2011.457

44. Oliva M, Demanelis K, Lu Y, et al. DNA methylation QTL mapping across diverse human tissues provides molecular links between genetic variation and complex traits. Nat Genet. 2023;55(1):112–122. doi:10.1038/s41588-022-01248-z

45. Hagberg AA, Schult DA, Swart PJ. Exploring Network Structure, Dynamics, and Function using NetworkX. Published online 2008.

46. Clauset A, Newman MEJ, Moore C. Finding community structure in very large networks. Phys Rev E. 2004;70(6):066111. doi:10.1103/PhysRevE.70.066111

47. Ashburner M, Ball CA, Blake JA, et al. Gene Ontology: tool for the unification of biology. Nat Genet. 2000;25(1):25–29. doi:10.1038/75556

48. Wu T, Hu E, Xu S, et al. clusterProfiler 4.0: A universal enrichment tool for interpreting omics data. The Innovation. 2021;2(3):100141. doi:10.1016/j.xinn.2021.100141

